# Shift in worker physiology and gene expression pattern from reproductive to diapause-like with colony age in the bumble bee *Bombus impatiens*

**DOI:** 10.1101/758367

**Authors:** Erin D. Treanore, Jacklyn M. Kiner, Mackenzie E. Kerner, Etya Amsalem

## Abstract

Insects maximize their fitness by exhibiting predictable and adaptive seasonal patterns in response to changing environmental conditions. These seasonal patterns are often expressed even when insects are kept in captivity, suggesting they are functionally and evolutionary important.

In this study we examined whether workers of the eusocial bumble bee *Bombus impatiens* maintained a seasonal signature when kept in captivity. We used an integrative approach and compared worker egg-laying, ovarian activation, body size and mass, lipid content in the fat body, cold tolerance and expression of genes related to cold tolerance, metabolism, and stress throughout colony development.

We found that bumble bee worker physiology and gene expression patterns shift from reproductive-like to diapause-like as the colony ages. Workers eclosing early in the colony cycle had increased egg-laying and ovarian activation, and reduced cold tolerance, body size, mass, and lipid content in the fat body, in line with a reproductive-like profile, while late-eclosing workers exhibited the opposite characteristics. Furthermore, expression patterns of genes associated with reproduction and diapause differed between early- and late-eclosing workers, partially following the physiological patterns.

We suggest that a seasonal signature, innate to individual workers, the queen or the colony is used by workers as a social cue determining the phenology of the colony and discuss possible implications for understanding reproductive division of labor in bumble bee colonies and the evolutionary divergence of female castes in the genus *Bombus*.

## Introduction

Insects maximize their fitness by exhibiting predictable and adaptive seasonal patterns in their physiology. Examples of the two most extensively studied phenomena are seasonal migratory movement and preparation for the winter-diapause. These phenomena allow an organism to survive unsuitable environmental conditions either through movement (migration) or avoidance (diapause) and are often characterized by shifts in metabolism, reproduction, and increased tolerance to environmental stressors (Kostal, 2006; Kostal and Denlinger, 2011; Ramenofsky and Wingfield, 2007). These adaptations evolved in response to changing environmental conditions, such as natural cycles of temperature, photoperiod, humidity and resource availability, leading to changes in insect abundance, mating, reproductive and dispersal habits as the season progresses (Denlinger, 2002; Gotthard, 2001). For example, decreased photoperiod and fluctuating temperatures can induce a reproductive diapause in the monarch butterfly (*Danaus plexippus)*, and reduced photoperiod leads to increased lipid accumulation in bean bugs (*Riptortus clavatus)* (Goehring and Oberhauser, 2002; Morita et al., 1999). Similarly, reduced resource availability induces dispersal by insects to areas more suitable for nest foundation and reproduction (Benton and Bowler, 2012) and reduced temperatures induce migratory behavior in butterflies and dragonflies (Chapman et al., 2015; Mikkola, 2003; Wikelski et al., 2006), overall allowing organisms to avoid an otherwise unsuitable environmental conditions and to synchronize their life history with other members of the population.

Insects often exhibit cyclic patterns associated with seasonality even when they are held in controlled conditions, provided with unlimited food or kept in captivity (Blake, 1960; Mao and Henderson, 2007; Miyazaki et al., 2014; Nisimura and Numata, 2003). Such a seasonal signature that is maintained regardless of the external conditions suggests functional importance, and highlights aspects related to the evolutionary history of species.

Social insects are a fascinating model in which to study these seasonal signatures in the wild and in the laboratory since seasonal changes may evolve to be used as social cues determining social behavior and the phenology of the colony. For example, honey bee (*Apis mellifera*) workers go through physiological changes throughout the season. Workers eclosing in the spring have reduced ovarian activation as compared to their summer and fall-eclosing sisters, likely a result of both reduced larval nutrition and cooler springtime temperatures (Hoover et al., 2006). Interestingly, honey bee workers exhibit these seasonal patterns even when reared in the laboratory under controlled conditions, with spring and fall workers possessing different nutritional requirements, aligning with the colony’s activities and needs across its yearly cycle (DeGrandi-Hoffman et al., 2018). Likewise, bumble bee workers (*Bombus terrestris*) eclosing in older laboratory colonies (comparable to late season colonies) show increased levels of constitutive immune defenses as compared to younger colonies (Moret and Schmid-Hempel, 2009). Ants provide several other fascinating examples: *Formica* ants reared in the laboratory under constant temperatures show distinct developmental patterns (Kipyatkov, 1993), and fire ant (*Solenopsis invicta*) workers from colonies collected in the summer and fall and maintained under laboratory conditions show distinct physiological phenotypes, with fall ants accumulating significantly more lipids (Cook et al., 2016). However, the extent to which the seasonal and the social cues are conflated and whether social cues are tied to innate seasonal signatures, that are independent from the surrounding conditions, is unknown.

In this study we examined whether workers of the eusocial bumble bee *Bombus impatiens* maintained a seasonal signature when kept in captivity. Bumble bees provide an excellent and tractable model system to examine this question for two reasons. First, bumble bees are annual eusocial species that go through a predictable life history trajectory as the colony ages. The newly mated queens disperse by late summer and enter a winter diapause lasting 6-9 months during which they are metabolically suppressed. Prior to diapause queens will mate, accumulate fat reserves such as lipids and their ovaries will remain inactive until the end of diapause. The transition to diapause is further characterized with a shift in hormone levels (e.g., Juvenile hormone) and gene expression associated with metabolism, stress and reproduction (Amsalem et al, 2015a). A single queen will then found a colony in early spring and the worker force is progressively built up to reach a size of several hundreds of workers by late summer (Amsalem et al., 2015b). The life cycle of the colony can be divided into two major social phases starting from the eclosion of the first worker (Duchateau and Velthuis, 1988). During the first phase (the pre-competition phase) that lasts approximately 4-5 weeks, the queen is the sole producer of eggs and workers function as helpers, whereas during the second phase (“competition phase”), workers may exhibit aggression and attempt to lay eggs, sexuals are produced and the social organization breaks down. While the triggers for the competition phase and the production of sexuals are not fully clear, it was suggested that workers modify their reproduction in response to changes in the queen (Almond et al., 2019; Amsalem et al., 2015a), the brood (Starkey et al., 2019a; Starkey et al., 2019b) and the demography of the colony (i.e., the number of workers) (Amsalem and Hefetz, 2011; Bloch, 1999; Van Doorn and Heringa, 1986). While earlier studies considered workers as a ‘clean slate’ (Amsalem and Hefetz, 2010), proposing that workers are morphologically, physiologically and behaviorally identical regardless of the time they eclosed during the life cycle, later studies show that workers eclosing at different times vary in intrinsic quality (Blacher et al., 2017; Couvillon et al., 2010; Shpigler et al., 2013). However, these investigations were limited to qualities such as body size and reproduction and did not explore further characteristics. Furthermore, it is unknown if these changes are driven by colony- or individual-level shifts in worker physiology.

Secondly, all bumble bees evolved from a eusocial species where all females were likely morphologically identical into a species where only large, fecund queens, but not workers, undergo a winter diapause (Cameron et al., 2007; Danforth et al., 2013; Hines, 2008; Wheeler, 1986). Bumble bee workers are, on average, three times smaller than the queens, do not mate, remain largely sterile and lack the physiological capacity to accumulate the fat body reserves necessary to survive the winter diapause (Amsalem et al., 2015b; Couvillon and Dornhaus, 2010). Thus, the differentiation of female castes could have been influenced by seasonal pressures shaping female physiology, an idea originally proposed for wasps where caste differences were suggested to be driven by nutritionally-sensitive ancestral genetic and physiological signatures that bifurcate during larval development in seasonal environments (the Diapause Ground Plan Hypothesis) (Hunt, 2006). This is further supported by the most recent reconstruction of the *Bombus* phylogeny suggesting that bumble bees were subjected to extensive period of cooling and are assumed to have radiated during the Miocene (Hines, 2008; Prokop et al., 2017), an epoch associated with gradual cooling that could have potentially shaped bumble bee social evolution. Overall, the maintenance and expression of a seasonal signature in females (both queens versus workers and within workers across the season) may generate predictions about the evolutionary trajectory of bumble bee caste divergence as a function of the environmental conditions.

To examine whether workers kept in captivity maintain a seasonal signature that corresponds to the age of the colony, we compared worker physiology and gene expression patterns throughout colony development. We hypothesized that bumble bee worker physiology changes as the colony ages and discuss different explanations underlying the observed changes. Specifically, we discuss four explanations for the physiological differences found in workers throughout the life cycle: (1) Worker physiology changes over time regardless of external conditions since they carry a signature indicative of their evolutionary history. Thus, the physiological signature is innate to workers. Here, we discuss The Diapause Ground Plan Hypothesis (Hunt, 2006) suggesting that social insect castes (sterile, short lived workers versus fecund long lived queens), diverged due to seasonal pressures; (2) Worker physiology is influenced by the social environment. Thus, the observed physiological changes are determined by the colony demography, can be controlled by changing the social environment and are not innate; (3) Workers evolved to use seasonal cues as social cues determining female reproduction. Thus, the observed physiological changes are innate to the colony; and (4) Worker physiology is dictated by the queen’s age at the time of egg-laying. Thus, the observed changes are innate to the queen.

To address the main question, we used an integrative approach and compared worker egg-laying, ovarian activation, body size and mass, lipid content in the fat body, cold tolerance and expression of genes related to cold tolerance, metabolism, and stress throughout colony development in colonies reared under controlled conditions. Genes were selected based on their role in physiological processes associated with seasonality, mainly diapause and reproduction. We examined the expression levels of (1) *foxo*: a forkhead transcriptional factor in the insulin pathway also shown to be down-regulated during diapause (Sim and Denlinger, 2008); (2) *menin*: a regulator of general stress response previously found to be up-regulated during diapause in *B. terrestris* queens (Amsalem et al., 2015a); (3) *hsp70:* a heat shock protein that is produced in response to cold in insects (Rinehart et al., 2007) and based on previous studies is expected to be up-regulated during diapause; (4) *pepck*: a metabolic enzyme involved in gluconeogenesis (Yang et al., 2009) that is expected to be down-regulated during diapause; (5) *hex1:* hexamerin storage protein that is produced in preparation for diapause (Hunt et al., 2007) and is expected to be up-regulated during diapause; (6) *mfe*: the final enzyme in the synthesis of Juvenile hormone (JH), the main gonadotropin in insects (Daimon and Shinoda, 2013) that is predicted to be downregulated during diapause and up-regulated in reproductives; and (7) *vitellogenin*: the major egg yolk protein female insects invest in the ovaries (Tufail et al., 2014) that is predicted to be down-regulated during diapause and up-regulated in reproductives.

## Materials and Methods

### Bumble bee rearing

*B. impatiens* colonies (n=13, Table S1 includes information on development and size of colonies used in this study, as well as the number of workers sampled from different colonies for each of the different analyses describe below) were obtained from BioBest at the age of 2-3 weeks after the eclosion of the first worker or reared in the laboratory from gynes produced in BioBest colonies using a standard CO_2_ treatment protocol. Briefly, newly mated queens are placed in a closed (but not sealed) plastic cage. A steady stream of pure CO_2_ is applied to the cage through a designated hole for 1 minute. Under these conditions CO_2_ reaches 100% within approximately 20 seconds (the time it takes bees to lose mobility) and remains this way for as long as the stream lasts. Queens are then kept in the cage for additional 30 minutes until they revived (Amsalem and Grozinger, 2017). Two more colonies were reared from queens caught in the wild in spring of 2018 in Newport and Landisville, Pennsylvania. All colonies were kept in constant darkness, 60% humidity and 28-30°C and manipulated under red light, that bees are unable to see. Bees were not allowed to leave the colony and pollen and 60% sucrose solution were provided *ad-libitum.* Bumble bees produce sexuals using a ‘bang bang’ strategy: an abrupt switch from the exclusive production of workers to the exclusive production of sexuals (Duchateau and Velthuis, 1988). Thus, in all experiments, workers were collected from queenright colonies (i.e., queen is present) throughout the natural development of the colonies until sexuals appeared. Workers eclosing early and late in the colony cycle (measured from the eclosion of the first worker) were considered early-and late-season workers, respectively.

### Bumble bee sampling

Sampling of workers from the parental colonies began at approximately two weeks after the eclosion of the first worker and continued until workers were no longer produced; colonies were queenright for the entire sampling period. Colonies were terminated when only males or gynes were exclusively produced for five consecutive days. Worker age was differentially controlled for in the different experiments. Age was not controlled for in the experiments examining ovary size, body mass and size, was partially documented in the cold tolerance experiment (in 2 out of 6 colonies) in order to verify our method of sampling (see below), was controlled for in egg laying assays (all workers were 10 days old), lipid content experiment (workers were 2-11 days old), and gene expression assays (all workers were 3 days old). In experiments where age was partially controlled or not controlled for, sampling occurred at a high-rate ensuring a constant turnover of individual workers in the colony and colony size of 20-30 workers at all times. To verify that our sampling method maintained a constant turnover of workers in the colony, we individual marked all workers (n=975) upon eclosion from a subset of the colonies used in our analyses and found that 92% of workers were sampled within a week of eclosion. We collected information about colony life span, the date of the production of first sexuals (males and gynes), the number of workers each colony produced, and which colonies were used in each analysis (Table S1). Overall, two colonies were used to examine worker body mass, size and ovary activation (n=472), four colonies were used to examine egg-laying in same-age workers (n=422), four colonies were used to examine fat body lipids (n=90), and six colonies were used to examine cold tolerance (n=2,417), and fat body gene expression (n=17).

### Ovary activation, body mass and size

To examine the reproductive potential of workers across the colony’s development (Table 1), groups of eight randomly sampled workers were removed from their parental queenright colonies every 2-4 days throughout the colony’s natural life cycle (weeks 3-13) and kept at −20° C until analysis. Age was not controlled for; however, sampling was done at a high-rate (the 2 colonies were sampled 59 times, 8 workers at a time, during a period of 10 weeks) ensuring a constant turnover of individual workers in the colony and a colony size of 20-30 workers at any time point. The length of the terminal oocyte in the three largest ovarioles was measured with a scaled ocular. *B. impatiens* females possess four ovarioles per ovary and at least one ovariole per ovary was measured. Mean terminal oocyte length for each bee was used as an index of ovarian activation. The same workers were measured for their wet body mass using an electronic scale and for their body size by measuring the width of the head as in Amsalem and Hefetz (2010).

**Table 1:** The list of examined variables and findings in this study

| VARIABLE |  | FUNCTION AND PREDICTIONS | RESULTS OF THIS STUDY |
| --- | --- | --- | --- |
| PHYSIOLOGY | Oocyte size | Females with a higher reproductive potential exhibit higher oocyte size | Smaller oocyte size in late-eclosing workers |
|  | Egg laying | Females with higher reproductive output lay more eggs | Lower number of eggs laid in late-eclosing workers |
|  | Cold tolerance | Females exhibit higher cold tolerance in preparation to diapause | Higher cold tolerance in late-eclosing workers |
|  | Body size | Females more adapted to diapause conditions are larger | Larger body size in late-eclosing workers |
|  | Body mass | Females more adapted to diapause conditions are heavier | Larger body mass in late-eclosing workers |
|  | Fat body lipid | Females more adapted to diapause conditions have higher fat storage | Higher fat storage in late-eclosing workers |
| GENE EXPRESSION | *vitellogenin* levels | Positively associated with female fertility. Reproductive females show higher *vg* expression compared with non-reproductives | No differences |
|  | *mfe* levels | Positively associated with female fertility. Reproductive females have higher *JH* levels compared with non-reproductives | Higher in early-eclosing workers |
|  | *hex1* levels | Encodes to storage proteins that accumulate in individuals during diapause | Higher in late-eclosing workers |
|  | *hsp70* levels | Encodes to heat shock protein that is higher in individuals during diapause | No differences |
|  | *menin* levels | A regulator of heat shock proteins and a general stress response, higher in individuals during diapause | Higher in early-eclosing workers |
|  | *foxo* levels | A transcriptional factor in the insulin pathway, lower during diapause | No differences |
|  | *pepck* levels | A gene coding to a metabolic enzyme used during gluconeogenesis. Reduced gluconeogenesis and a conservation of energy stores is typical to individuals during diapause | Higher in early-eclosing workers |

### Egg laying

To examine the physiological aptitude of queen-less workers (i.e. queen is absent) to lay eggs, we counted the number of eggs laid by same-age (10-day old) workers sampled from four colonies throughout the entire life cycle (weeks 2-10). Two newly-eclosed workers of similar size were removed from their parental colony and placed into a cage (11 cm diameter x 7 cm tall) where they were provided *ad libitum* pollen and 60% sucrose syrup. Cages were inspected for the total number of eggs laid by workers at the end of the 10-day period. Post analysis, the data were divided to three time points (earlier than week 5, weeks 6-8 and week 9 or later), based on the division observed in the ovary experiment where these groups clustered together. These timepoint also matched the life cycle phases of *B. impatiens* (pre competition, post competition, and sexual production).

### Cold tolerance

To examine cold tolerance of workers across the colony’s development, we used a modified protocol from Karsai and Hunt (2002). Groups of 10 workers were randomly sampled from their parental colony throughout the entire colony cycle (weeks 3-13). In two colonies all workers were individually marked daily upon eclosion and put back into the colony, making up a total of 40% of all workers analyzed for cold tolerance. Workers that had their age documented were 2-21 days old at the time of sampling and 92% of them were sampled within 7 days of eclosion. Workers were placed in a 28 × 22 × 5 cm cardboard box and into a refrigerator mimicking diapause-like conditions (70% humidity, total darkness, and 4°C). Sampling frequency varied depending on colony growth, but a minimum of 10 workers were sampled from every colony for every week of the experiment. To assess survival of workers, boxes were removed from the fridge every 48 hours and were exposed to ambient light and temperature for ten minutes. Workers were lightly prodded with forceps to verify whether they were alive and any that showed no sign of movement in 10 minutes were considered dead and stored at −20° C until further analyses. Live workers were placed back into the fridge. In order to identify potential confounding factors leading to mortality in cold, other than the colony age, we examined the correlation between survival length and worker age, body mass and oocyte size in a subset of the dead workers (Table S2).

### Lipid analyses

To examine worker fat body lipid content, we compared lipid amounts in early-season (weeks 3-5) and late-season (weeks 8-11) workers, taken from four colonies. Workers were individually marked upon eclosion and worker age upon sampling was recorded (2-11 days). To control for size variation, worker body and eviscerated abdomen mass were measured prior to dissections. Lipid analysis was done as in Amsalem (2015a) and Amsalem and Grozinger (2017). Briefly, the abdomen was separated from the thorax and eviscerated of all content except the fat body (i.e., the ovaries, gut and sting complex were removed), and placed in 2% sodium sulphate. Two hundred microliters of the fat body homogenate were added to glass vials containing 2.8 mL chloroform/methanol mix. Vials were centrifuged (x3000g) and the supernatant was mixed with 2 mL of distilled H_2_O separating between the upper aqueous fraction and the lower organic fraction (lipids). The lower organic lipid fraction was separated and quantified using a vanillin-phosphoric acid reaction. A standard curve for lipids was created using five different concentrations of 0.1% vegetable oil diluted in chloroform. Absorbance values (OD 525) for each sample were measured with a Nanodrop One™ and converted to micrograms per worker, based on a formula calculated from the regression line derived from the calibration curve. The amounts of lipids are presented as percentages of the abdomen mass to correct for body size.

### Gene expression

Expression of genes was compared in 3-day old workers sampled from early season colonies during weeks 2-4 (8 workers taken from 3 colonies) and from late-season colonies during weeks 8-10 (9 workers taken from 3 colonies; Table S1). Expression levels of *foxo, hsp70, hex1, menin, vg, mfe*, and *pepck* were examined in the workers’ fat body. Target genes were selected based on their biological relevance to cold tolerance, lipid metabolism, stress and reproduction (Amsalem et al., 2015a; Hunt et al., 2007; Jedlicka et al., 2016; Spacht et al., 2018). Design of forward and reverse primers for each gene was done using the primerBLAST (https://www.ncbi.nlm.nih.gov/tools/primer-blast/) and are provided in Table S3. Primers were designed to target the most conserved region between sequences.

Workers were individually marked in their parental colonies upon eclosion, flash-frozen 72 hours later and stored at −80° C until use. To isolate the fat body tissue, workers were dissected using RNAlater to separate the abdomen from the thorax and remove all abdominal contents, including ovary tissue, sting complex, and the gut. The remaining carcass and attached fat body were immediately placed in lysis buffer and stored at −80° C. Total RNA samples were extracted using the RNAeasy mini kit (Qiagen, Valencia, CA, USA) according to the manufacture instructions. cDNA was synthesized following the manufacturer’s instructions using 200 ng of RNA (Applied Biosystems).

Gene expression was measured on a QuantStudio 5. For real time PCR reactions, 2 μL of diluted cDNA were combined with 5 μL SYBR Green (Bioline, Luckenwalde, Germany), 0.2 μL of each forward and reverse primer and 4.6 μL DEPC-water. To control for PCR efficiency and individual differences across samples, we used two reference genes: *Arginine kinase* and *Phospholipase A2*. These genes have been previously used in several studies with *B. impatiens* (Amsalem et al., 2015a; Amsalem and Grozinger, 2017). Reactions were performed in triplicate for each of the samples and averaged for use in statistical analyses. Negative control samples and a water control were present on each plate. Expression levels of candidate genes were normalized to the geometric mean of the two housekeeping genes using the 2^−ΔCt^ technique (Vandesompele et al., 2002). For visual presentation, expression levels were normalized to the treatment with the lowest expression level.

### Statistical analyses

All statistical analyses were conducted using JMP Pro. software v.14.0.0 (SAS Institute Inc., Cary, NC, USA). Data were tested for normality using Goodness of fit test prior to the statistical analysis and were analyzed using an ANOVA mixed model or a generalized linear model (GLM), accordingly. The effect of parental colony was examined and included in the model if found significant. Oocyte size data were analyzed using a generalized linear model (GLM) with colony age as the fixed effect, Poisson distribution, and log as the link function. Egg laying and worker size data were analyzed using an ANOVA mixed model with parental colony as a random effect followed by a post-hoc analysis using Tukey’s HSD Test. Worker mass was analyzed using a GLM with colony age as the fixed effect, a normal distribution, and log as the link function. Cold tolerance data were examined using Kaplan-Meier survival analyses. Lipid data were normally distributed following a log-transformation and analyzed using an ANOVA mixed model with parental colony as a random effect. Gene expression data were examined using either a t-test or a Wilcoxon ranked test, depending on the distribution of the data.

## Results

### The effect of colony age on worker oocyte size and egg laying

Worker oocyte size (Figure 1) was significantly affected by colony age (GLM, χ^2^_10_=85.88, p<0.001) but not by the parental colony (GLM, χ^2^_1_=1.64, p=0.20). The data clustered to three time points, separating between early season (<5 weeks), mid-season (6-9 weeks) and late season (>9 weeks). Post-hoc contrast tests were performed to compare early, mid, and late-season workers. Early-season workers (weeks 3-5) had significantly smaller oocyte size compared with mid-season (6-9 weeks) workers (0.31±0.04 mm in early vs. 0.83±0.06 mm in mid-season, means ± SEM), in line with the pre-competition and competition phases of the colony (Duchateau and Velthuis, 1989). However, late-season workers (weeks 10-12) also showed smaller oocyte size (0.36±0.03 mm) compared with mid-season, despite the inability of the queen to inhibit reproduction at this stage (Duchateau and Velthuis, 1988; Duchateau and Velthuis, 1989). The effect of colony age on worker oocyte size was the same regardless if worker body mass or worker size were included in the model.

**Figure 1:**
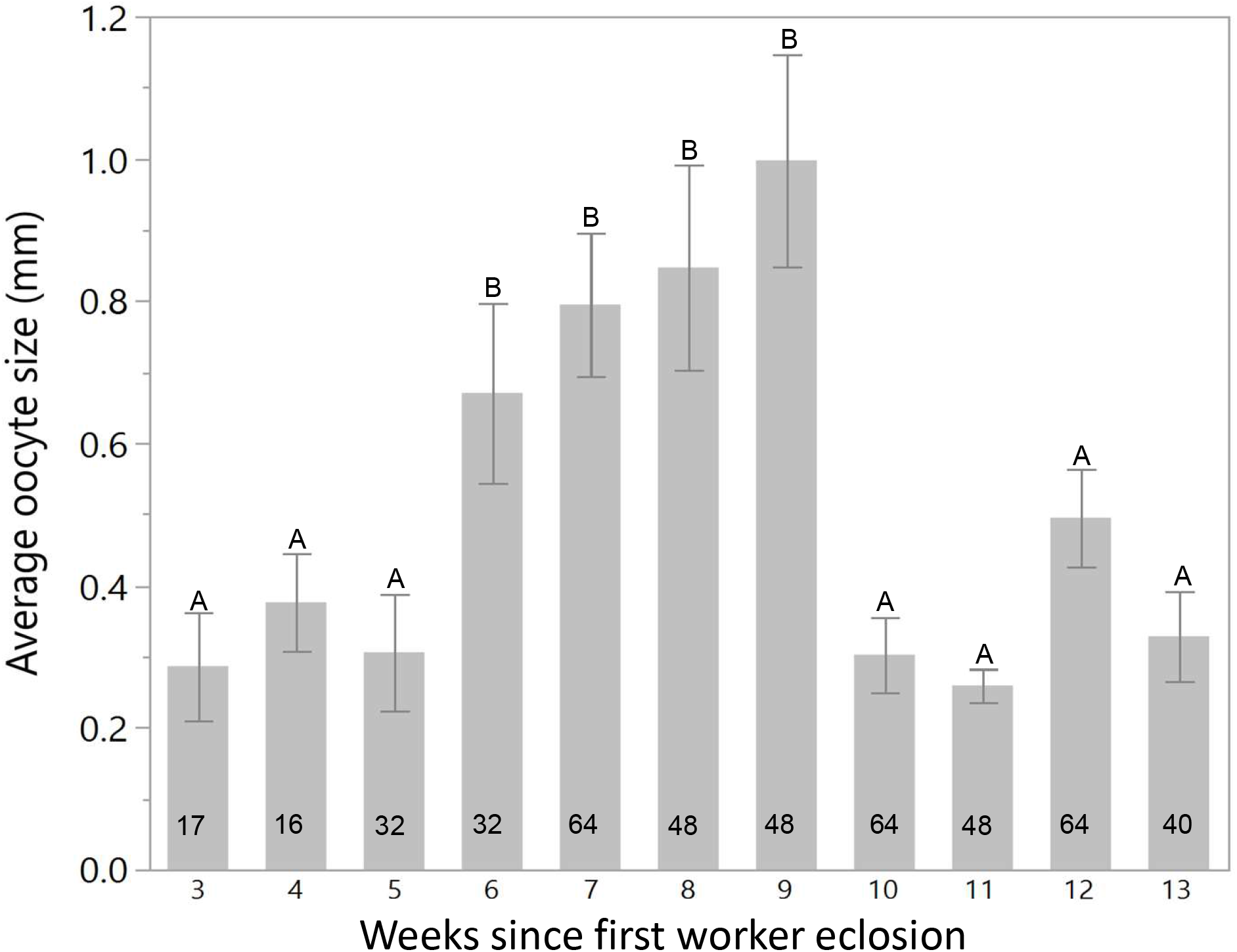
The effect of colony age on *B. impatiens* worker oocyte size. Workers were sampled from two colonies during weeks 3-13 (calculated from the eclosion of the first worker) and immediately frozen (n=472 workers). Numbers in columns represent the number of individuals and letters in bars denote statistical differences at α=0.05. Data are presented as means ± SE.

Since the presence of the queen may mask the differences in worker reproduction early in the colony’s life cycle, we also examined the effect of colony age on the physiological aptitude of workers to lay eggs in queen-less conditions (Figure 2). We placed pairs of newly eclosed workers in plastic cages for 10 days (n=422 workers) and compared their cumulative egg laying. Workers were collected from three time points during colony development, based on the findings in the previous experiment (weeks 2-5, 6-8 and >9 weeks) and the overall longevity of the colonies used for these experiment (colonies were terminated at the age of 9-10 weeks, thus week 9 was considered late season). The average number of eggs was significantly affected by colony age (f_2,206_= 6.35, p=0.002). On average, workers that eclosed after week nine laid significantly fewer eggs (16.77 ± 1.5) compared to the number of eggs laid by workers that eclosed during weeks 2-5 (23.4±1.32) or weeks 6-8 (25.41±1.42). The parental colony did not significantly affect the number of eggs laid by workers (f_2,206_= 4.51, p=0.23), and thus was not included in the model.

**Figure 2:**
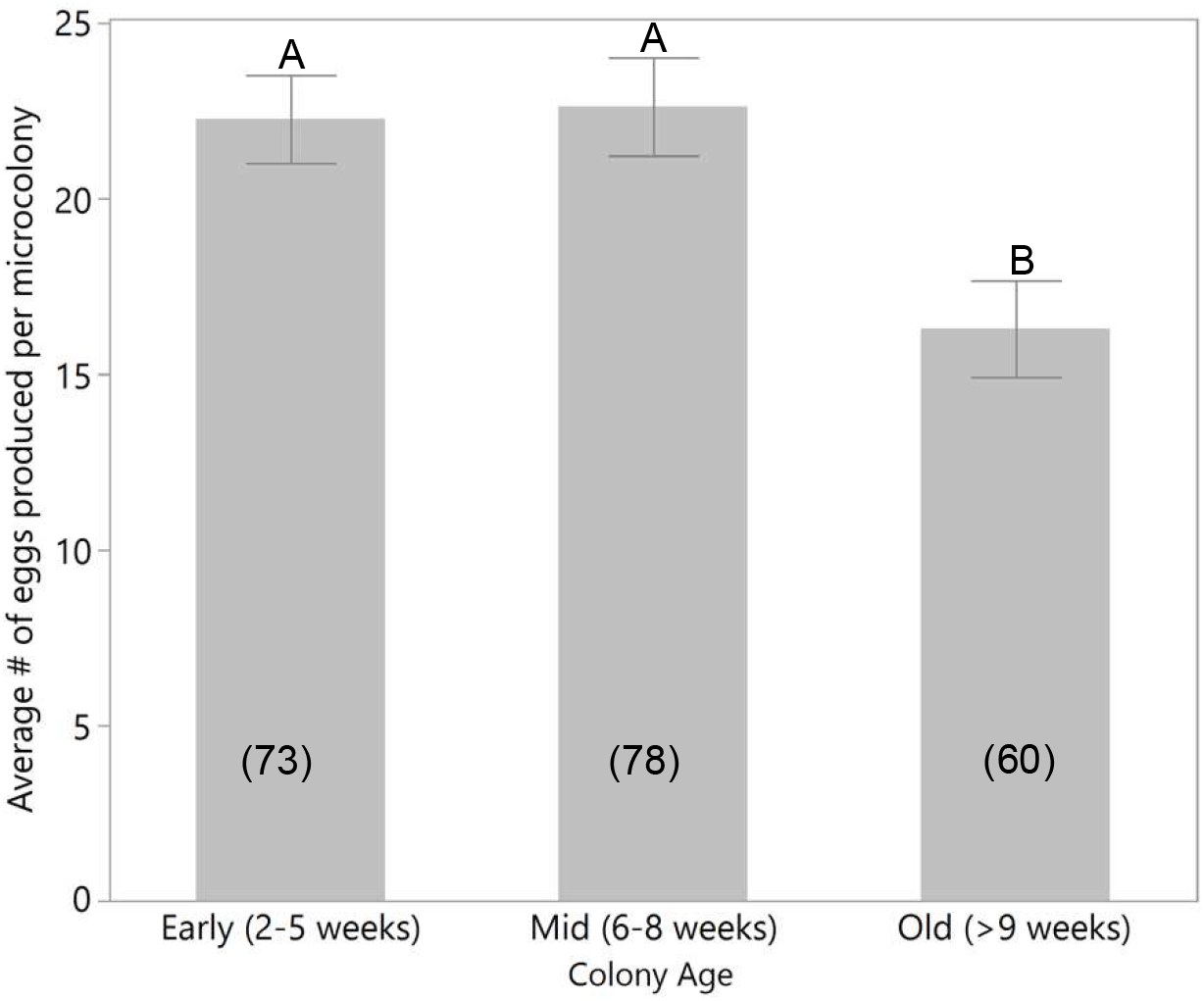
The effect of colony age on the average cumulative number of eggs laid by queen-less *B. impatiens* workers. Newly eclosed workers were sampled from four colonies during weeks 2-11 (calculated from the eclosion of the first worker) and kept in pairs for 10 days (n=211 cages). Numbers in columns represent the number of cages and letters above bars denote statistical differences at α=0.05. Data are presented as means ± SE.

### The effect of colony age on workers cold survival

On average, workers survived 11 days under diapause-like conditions (3-5°C, 70% humidity) (Figure 3). Colony age (in weeks) significantly affected the number of days workers survived in diapause-like conditions (Kaplan-Meier log-rank test χ^2^_10_ =81.5, p<0.001), with a difference of approximately three days between the average survival in the earliest week (week 3, 10.05±0.45 days) compared with the latest week (week 13, 13.15±0.50 days). Significant differences were also found in the average survival of workers from wild and commercial colonies, with workers from wild-caught colonies surviving on average 11.59±0.2 days while workers from commercial colonies survived on average 10.8±0.1 days (Kaplan-Meier log-rank test, χ^2^_1_=17.34, p<0.001). The parental colony had a significant effect on worker survival in both wild (Kaplan-Meier log-rank test, χ^2^_1_=9.85, p<0.001) and commercial colonies (Kaplan-Meier log-rank test, χ^2^_3_=133.57, p<0.001), with a difference of three days on average between the colonies that survived the longest and the shortest.

**Figure 3:**
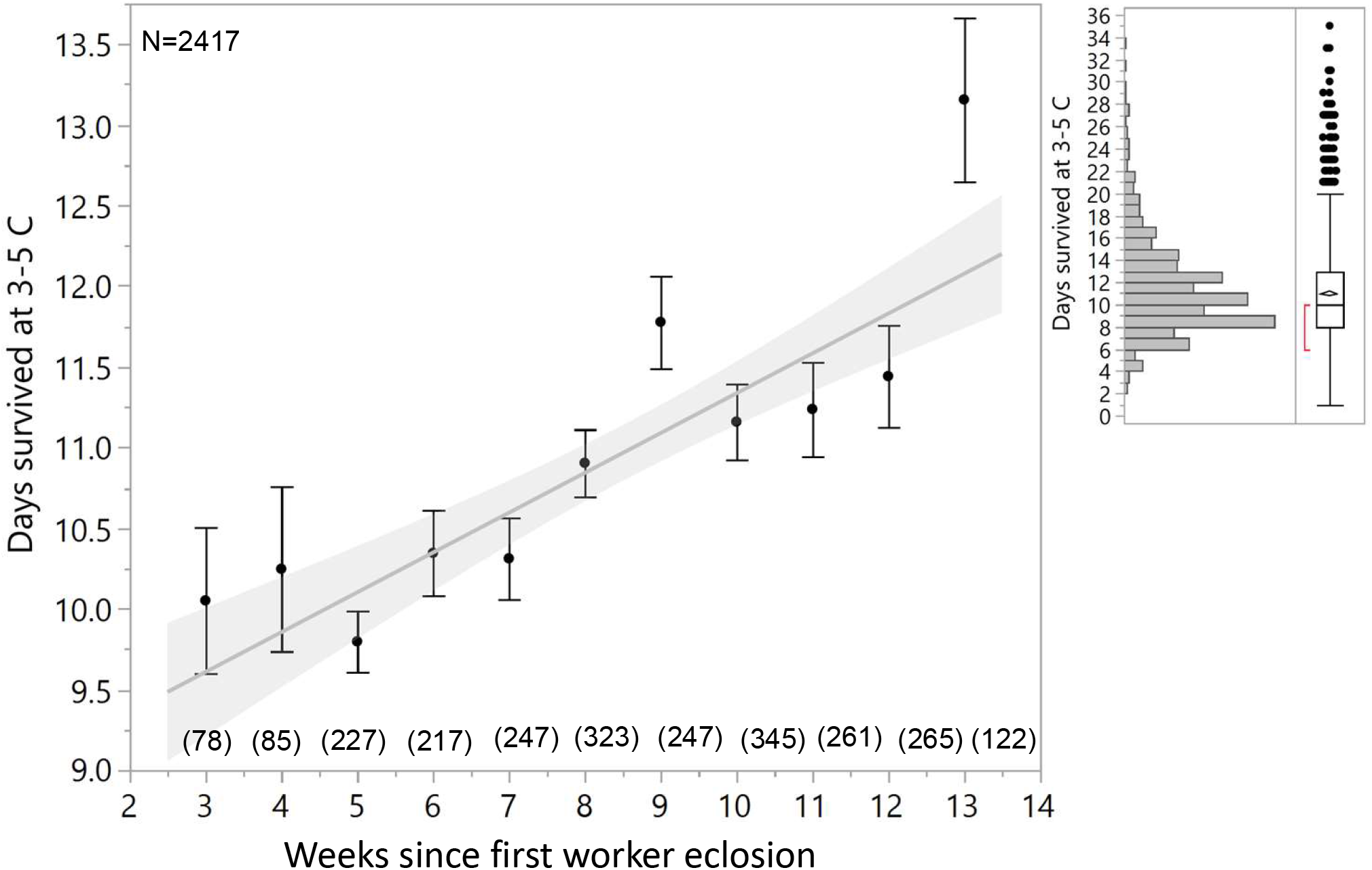
The effect of colony age on *B. impatiens* worker survival in diapause-like conditions (3-5°C, 70% humidity). Group of 10 workers were sampled every few days during weeks 3-13 from six colonies (four commercial and two wild-caught, n=2,417 workers). The figure to the right represents the distribution of the number days workers survived in cold in the total sample size. Numbers in brackets represent the total number of workers sampled per week. Statistical significance was accepted at α=0.05. Data are presented as means ± SE.

To test whether mortality in cold could be attributed to factors other than colony age, we examined the correlations between the number of days survived in cold and several potential confounding factors. Worker’s age was documented in two colonies where all workers were individually marked upon eclosion. We found a significant, though weak, negative correlation between worker age and cold survival in both colonies (Colony 3: Pearson’s r= −0.14, n=232, p=0.03; Colony 4: Pearson’s = −0.21, n=743, p<0.001). The body mass of workers upon death was examined in all six colonies but was weakly positively correlated with survival only in two colonies (Colony 1: Pearson’s r=0.13, n=322, p=0.02; Colony 3: Pearson’s r=0.22, n=233, p<0.001). Finally, no significant correlation was found between oocyte size upon death and survival in any of the six colonies (Table S2).

### The effect of colony age on worker body mass and size

There was a significant effect of colony age on worker size (f_1,471_=3.89, p=0.05) and worker body mass (GLM, χ^2^_10_=125.19, p<0.001; Figures 4a and 4b, respectively), though the size effect was slightly weaker than the mass effect. There was an increase in both size and mass over time, up to week 12, and a slight decrease in the last week (week 13). Colonies typically start showing signs of collapse after weeks 9-10 (see also Table S1 for the average survival in weeks of all colonies). Thus, we attributed the decrease in mass/size in week 13 to the final stage of the colonies and believe the consistent trend of increasing body size and mass in weeks 2-12 reflect the colony’s normal development more accurately than the last week. Neither body mass (GLM, χ^2^_1_=2.76, p=0.10) or size were affected by the parental colony (f_1,470_=4.03, p=0.84).

**Figure 4:**
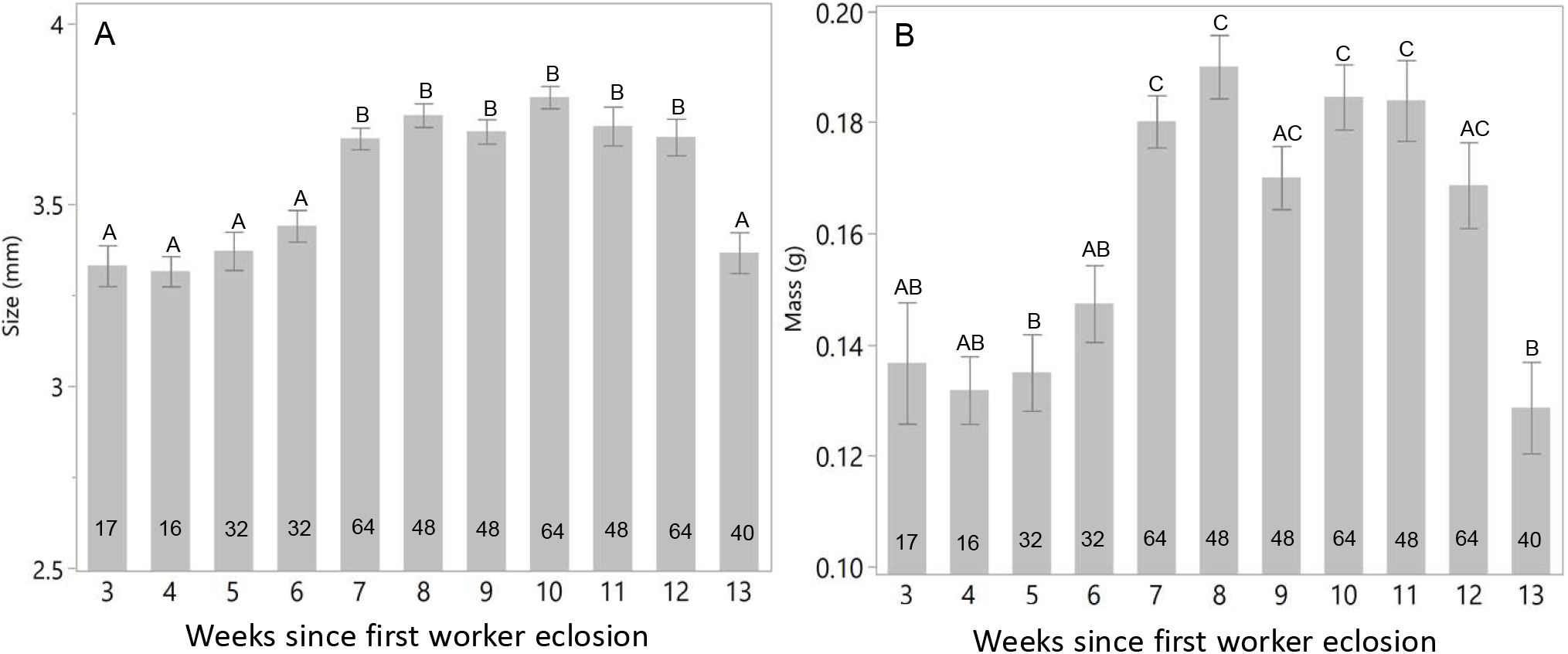
The effect of colony age on *B. impatiens* worker head width size (A) and body-mass (B). Workers were randomly sampled from two colonies during weeks 3-13 (calculated from the eclosion of the first worker, n=472 workers). Numbers in columns represent the number of cages and letters above bars denote statistical differences at α=0. Data are presented as means ± SE.

### The effect of colony age on worker fat body lipids

Lipid content in the fat body (normalized to eviscerated abdomen mass) was significantly higher in late-season compared with early-season workers (ANOVA Mixed Model: f_1,85.1_=5.41, p=0.023, Figure 5). Lipid content was not affected by the parental colony (f_1,85.1_=5.41, p=0.33), and no significant correlation was found between worker age and fat body lipid content (Pearson’s r=0.08, n=90, p=0.42).

**Figure 5:**
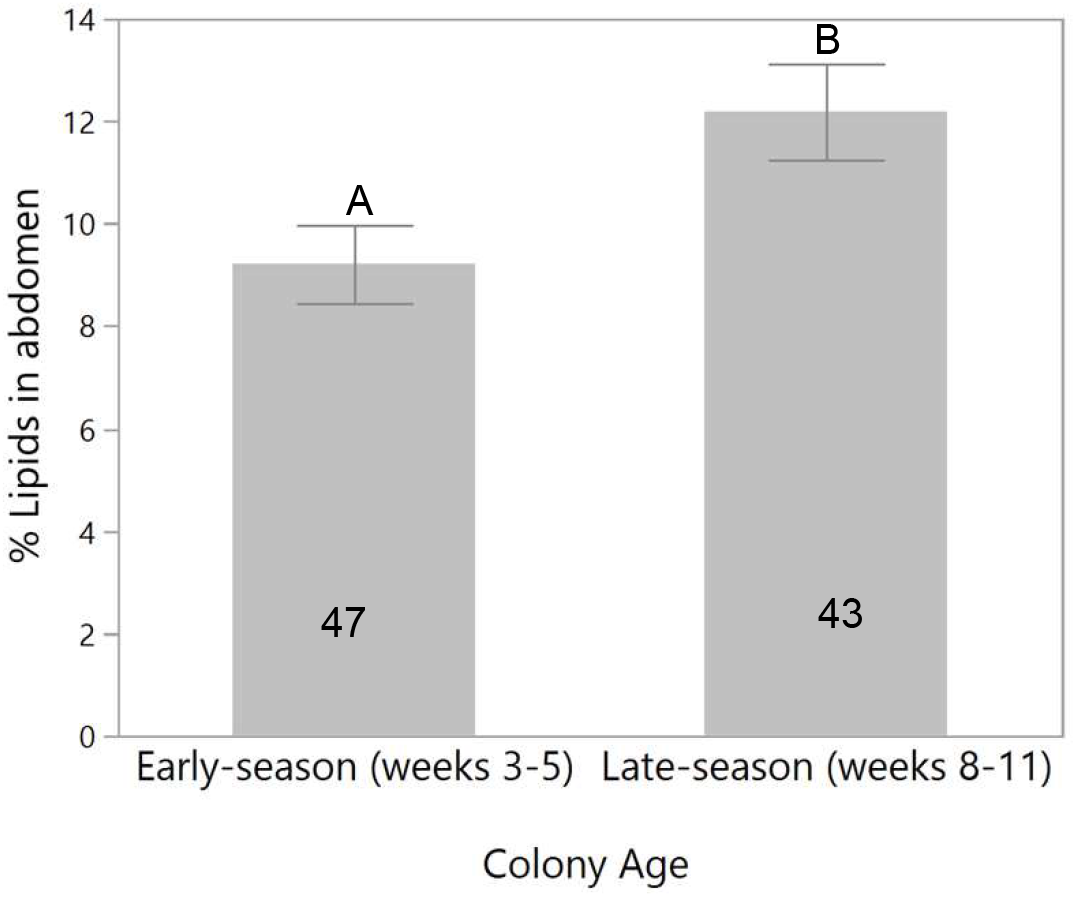
The effect of colony age on the average percentage of fat body lipids in *B. impatiens* worker’s abdomens. Workers (n=90 workers) were sampled from four colonies during weeks 3-5 (early-eclosing) versus weeks 8-11 (late-eclosing). Numbers in columns represent the numbers of individuals and different letters above bars denote statistical significance at α=0.05. Data are presented as means ± SE.

### The effect of colony age on worker gene expression

Expression levels of selected genes were examined in 3-day old workers (Figure 6). We chose 3-day old workers since they are too young to activate their ovaries, ensuring that gene expression differences are not the result of a different reproductive state. Gene expression levels were compared using a t-test or Wilcoxon ranked test, depending on whether the data were distributed normally or not (see Methods). Expression levels of *menin, mfe* and *pepck* were significantly higher in early-compared with late-season same-age workers (*menin*: *t*-test: t(10.88)=2.79, p=0.008; *mfe*: Wilcoxon ranked test: χ^2^_1_=5.36, p=0.02; *pepck*: *t*-test: t(12.16)=2.25, p=0.02). Expression levels of *hex1* were significantly higher (Wilcoxon ranked test: χ^2^_1_=3.57, p=0.05) and expression levels of *hsp70* were marginally insignificantly higher (*t*-test: t(8.67)=1.5, p=0.08) in late- vs. early-season workers. The effect size found in our study, although not very high (typically 2-3 times higher in significantly upregulated genes), is equivalent to findings in previous studies. For example, in Amsalem et al, (2015a) where we compared the expression of *foxo* in diapausing vs. non diapausing queens in *B. terrestris*, the expression levels were 2 times higher during diapause. *Foxo* is a well conserved gene known to be upregulated during diapause, so the size effect in this study for most of the genes is not out of norm. No significant differences were found in the expression levels of *vg* (χ^2^_1_=1.12, p=0.29) or *foxo* (χ^2^_1_=2.08, p=0.15).

**Figure 6:**
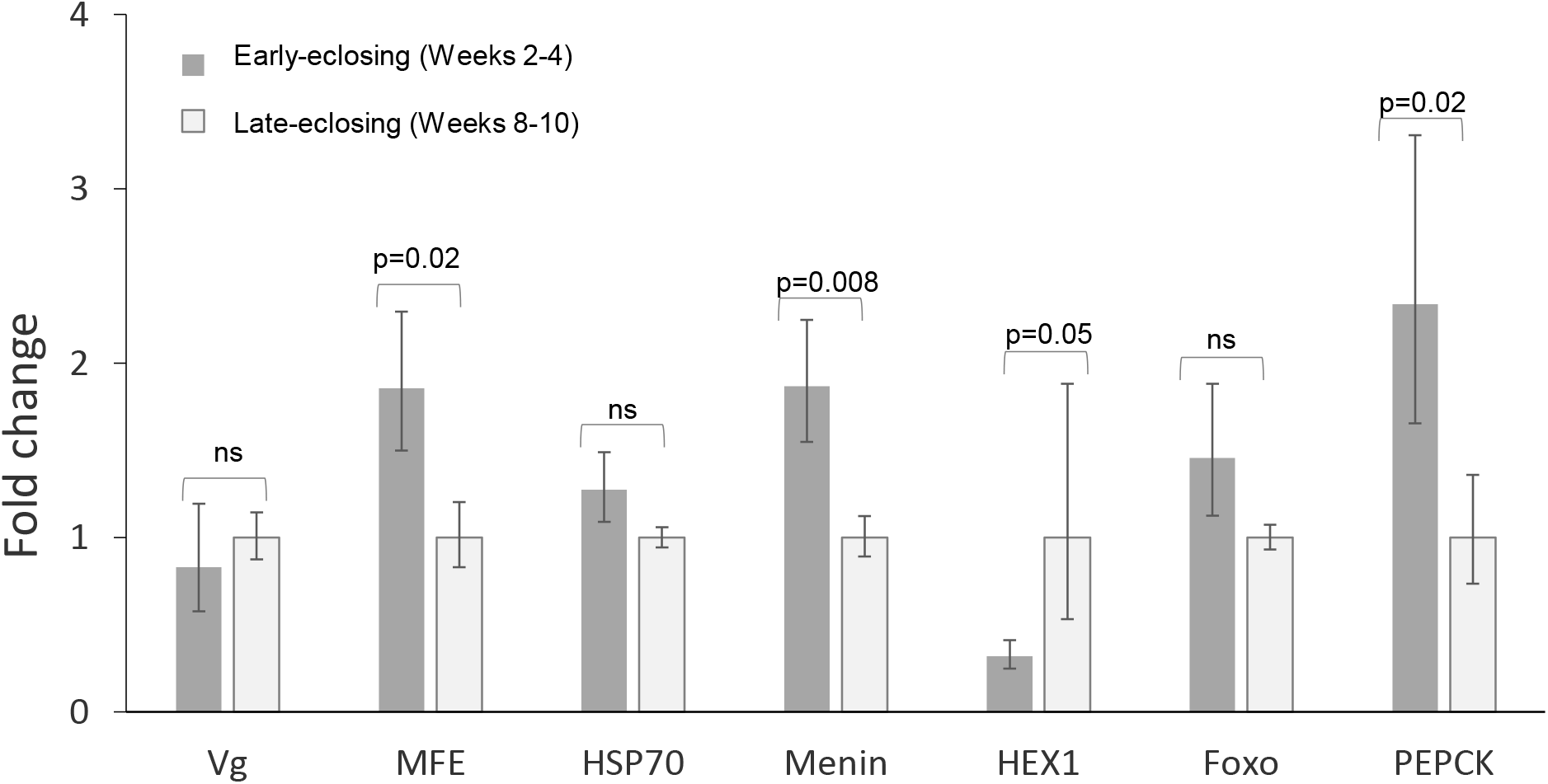
The effect of colony age on the expression levels of several candidate genes in 3-day old *B. impatiens* workers. Workers were sampled from 6 colonies during weeks 2 and 10 (n=17 workers). Different values above bars denote statistical significance at α=0.05. Data are presented as means ± SE.

## Discussion

Our study demonstrates that bumble bee workers exhibit physiological and gene expression pattern shifts from reproductive to a diapause-like profile as the colony ages, despite being reared under controlled temperature and humidity and provided unlimited food. Overall, workers eclosing late in the colony cycle had less activated ovaries under queen-right conditions and laid fewer eggs under queen-less conditions compared with workers eclosing early in the life cycle. On average, late-eclosing workers were larger, heavier, stored more lipids and were able to tolerate cold longer compared with early-eclosing workers. These observations are in line with an overall reproductive-like physiology early in the life cycle and a diapause-like physiology late in the life cycle.

These results are also summarized in Table 1. The main question to be answered is why maintaining and expressing these signatures is adaptive for bumble bee workers. Although not mutually exclusive, below we discuss four possible explanations for this phenomenon.

### 1. Physiological shift in workers is innate to workers: Late-season worker physiology reflects an ancient diapause signature

Circannual patterns regulating developmental and physiological processes in individual insects, although uncommon, have been documented across numerous taxa (Miyazaki et al., 2014; Saunders, 2009). Perhaps the most well-known example is the carpet beetle (*Anthrenus verbasci),* whose larvae possess an endogenous annual rhythm that controls the onset of diapause (Blake, 1960; Nisimura and Numata, 2003). Dragonflies also use a circannual rhythm to alternate between phases of diapause and development over the course of a year (Norling, 1984; Saunders, 2010). In social insects, termites (*Coptotermes formosanus*) were shown to a have circannual rhythm regulating the developmental time of the soldier caste (Mao and Henderson, 2007). As circannual patterns allow an insect to synchronize their activities, physiology, and behavior with annual environmental fluctuations, its adaptive value may be particularly important in seasonal environments where insects must survive unfavorable periods.

In species that inhabit seasonal environments, larger females are better suited to survive the winter-diapause, a physiologically demanding period during their life cycle, and found a colony the following year. Selection based on body-size can lead to the divergence of female castes and to fixed morphological differences between longer-lived, larger, diapause-capable queens and shorter-lived, smaller, helper workers that are unable to survive the winter. Our finding that late-eclosing workers exhibit a more diapause-like profile compared to early-eclosing workers is similar to solitary multivoltine bee species (e.g., several Xylocopa species) whereby the early-season generation enter the reproductive cycle immediately following eclosion, and the late-season generation refrain from reproduction and enter diapause. This may suggest that the shift from immediate reproduction (‘reproduce now’) to diapause (‘reproduce later’) played an important role in the divergence of female castes in bumble bees, an idea that was formulated as the Diapause Ground Plan Hypothesis (DGPH) (Hunt, 2006).

The DGPH proposes that caste differences are driven by nutritionally-sensitive ancestral genetic and physiological signatures that bifurcate during larval development in seasonal environments, resulting in two trajectories: (1) The ‘G1 pathway’ where early-season, poorly-fed larvae develop slowly into adults with low-fat and protein stores exhibiting a reproductive-like physiology. These adults are predicted to seek protein upon eclosion and attempt to reproduce right away (‘reproduce now’); (2) The ‘G2 pathway’ where late-season, well-fed larvae develop faster into adults with high-fat and protein stores and exhibiting a diapause-like physiology that allows them to postpone reproduction until the following year (‘reproduce later’). These two trajectories are hypothesized to give rise to a solitary ancestor with a facultative physiological and genetic signature from which small females, incapable of entering diapause (workers) and large, diapause-capable females (queens) have evolved.

Empirical support for the DGPH in insects is scarce and mostly indirect. Gyne-destined larvae of *Polistes* wasps (potential future queens) carry a diapause-signature, exhibiting a longer developmental period and higher levels of hexamerin storage proteins that are associated with preparation for diapause (Hunt et al., 2007). Study in *B. terrestris* showed that while the physiological and molecular changes associated with diapause are well-conserved across species (Amsalem et al., 2015a; Ragland and Keep, 2017), genetic pathways regulating diapause in social insects have been co-opted to regulate the divergence of castes, evidenced by substantial overlap between genes related to caste determination and diapause in bumble bees, and a shift in the role of key genes regulating diapause in *B. terrestris* compared with Diptera species (Amsalem et al., 2015a; Ragland et al., 2010). Finally, a recent phylogeny demonstrated that the most recent common ancestor of all bees likely exhibited developmental diapause, and that an ancestral shift from developmental diapause to adult diapause was an important preadaptation to sociality in bees (Santos et al., 2019).

Bumble bee female castes are fixed and are determined during larval development, leading to queens that are three time larger than workers (Couvillon and Dornhaus, 2010). These queens, unlike workers, are capable of surviving winter diapause and consequently of initiating a colony in the next spring. Our study did not directly examine the differences between queens and workers, but may provide insights into the evolutionary history of bumble bee castes by examining the shift in the seasonal signature workers expressed. Bumble bee workers show large variability in body size (Couvillon and Dornhaus, 2010), which increases toward the end of the season (Knee and Medler, 1965; Shpigler et al., 2013), and also exhibit a shift between reproductive to diapause-like profile as the colony ages. These correlations may suggest that co-option of pathways regulating diapause to generate reproductive and non-reproductive phenotypes may have facilitated the evolution of the worker caste in species that evolved in seasonal environments. In support of that, late-eclosing workers in our study were larger and heavier (Fig. 4), and also had higher amounts of lipids in their fat body (Fig. 5), both of which are characteristic of the diapause phenotype. They also showed higher cold tolerance (Fig. 3) and lower reproductive potential (Fig. 1-2). Furthermore, genes that are predicted to be altered in preparation to diapause were differentially regulated in early vs. late-eclosing workers, in line with the DGPH (Fig. 6).

Environmental factors affecting caste divergence have been demonstrated in several species. For example, the production of discrete phenotypes in ants of temperate climates is influenced by interaction between genetics and temperature (Libbrecht et al., 2013; Schwander and Keller, 2008), and the effect of nutritional cues during development on caste determination was demonstrated in bees, wasps, ants and termites (Corona et al., 2016). These examples and the data presented in this study suggest that the DGPH warrants further direct investigation. Future studies examining how seasonality affects larval development and body size across social species and manipulation of nutritional cues in larvae could further elucidate whether the diapause ground plan was a prerequisite for the development of reproductive and non-reproductive castes. A large-scale examination of genes that vary with seasonality in comparison with genes regulating diapause across species will further elucidate how conserved these pathways are and the role diapause may have played in the evolution of sociality and castes.

### 2. Physiological shift in workers is not innate: Worker physiology is influenced by the social environment

The observed shift in worker physiology could also be a result of the social environment during larval development (i.e., the number of workers, the amount of food fed to the larvae and the presence of older nestmates). However, we reject this explanation for several reasons. In several of our experiments (Fig. 1-5) we didn’t sample workers before weeks 2-3 (i.e., until sufficient number of workers has accumulated) and controlled for the number of workers in the colony at the time of sampling, maintaining no more than 20-30 workers and the queen throughout the colony cycle. Therefore, the larger and heavier workers produced by the end of life cycle were not taken care of by a higher number of individuals during their larval development. It should be noted, however, that a previous study found that *B. terrestris* worker body size was influenced by the social environment, with smaller-size workers produced when reared by one versus ten queen-less workers (Shpigler et al., 2013). However, in the same study, no such difference was found under queen-right conditions that better reflect our experimental design: workers that were reared by the queen and >10 workers were not larger than workers reared by a single worker in two out of three replicates (Shpigler et al., 2013). Interestingly, the sheer number of workers in late-stage colonies was proposed as a possible explanation for the decrease in queen dominance and the initiation of worker reproduction (Van Doorn and Heringa, 1986). However, this was not supported by our data showing that the shift in worker ovarian activation occurred despite the controlled colony size.

A previous study conducted in *B. terrestris* showed a pattern of workers ovary activation throughout colony development similar to the one we found (Fig. 1), with lower ovary activation in workers in both very early and very late in the colony cycle (Bloch and Hefetz, 1999). The authors suggested that early inhibition of worker reproduction is induced by the queen while the later inhibition is caused by nestmate workers. This explanation, too, does not line up with our data since the frequent sampling of workers likely prevented the formation of such a hierarchy among workers.

Lastly, in all experiments we provided the bees with unlimited food of the same quality, eliminating the possibility that fluctuations in resource abundance or quality can explain the differences we observed. All colonies and individuals were kept in darkness, 60% humidity and 28-30° C, and overall, despite the control of worker demography, external conditions and queen presence we still found clear differences in the physiology of early- and late-eclosing workers. However, our data are correlative in nature as they were collected from normally developing colonies. Thus, we cannot preclude the potential effect of other internal factors (e.g., brood number, brood composition, queen age, etc.) that were not controlled for.

### 3. Physiological shift in workers is innate to the colony: Workers evolved to use seasonal cues as social cues determining female reproduction

A bumble bee colony typically contains hundreds of workers that emerge progressively over a period of approximately three months, followed by dispersal of reproductives and dissolution of the colony (Michener, 1974). In nature, this period is characterized by progressive decreases in floral resource availability (pollen and nectar), ambient temperatures, and photoperiod. During the first phase of the colony cycle the queen exerts full control over worker reproduction but later in the season workers start activating their ovaries and laying eggs (Amsalem et al., 2015a; Duchateau and Velthuis, 1988). Workers are unmated, thus can only produce haploid eggs that develop into males and depend on the queen to produce the worker force to support the rearing of sexuals and to switch into rearing gynes, two events that are controlled solely by the queen (Cnaani et al., 2000; Duchateau and Velthuis, 1988). In *B. terrestris* it was demonstrated that workers ‘eavesdrop’ on the queen by possibly monitoring her age and reproductive output and waiting for her to lay eggs that develop into gynes before attempting to lay eggs (Alaux et al., 2006). Workers further closely monitor the queen’s fecundity and initiate aggressiveness towards her when her productivity decreases (Almond et al., 2019), often resulting in matricide (Bourke, 1994). Therefore, it is beneficial for workers to be physiologically prepared to lay eggs in the event that the queen dies prematurely.

Seasonal limitations dictate that if workers wait too long to reproduce, even if successful in laying eggs, their eggs are unlikely to complete development due to decreased floral resources and temperatures. Furthermore, even in the case that eggs are fully developed to adults, the eclosion of males that were laid too late in the season is unlikely to synchronize with the eclosion of gynes resulting in inability to find a mate. Therefore, while investing in reproduction early in the season is beneficial, investing in reproduction late in the season is likely to be futile.

In accordance with this explanation, we found that workers exhibit a higher reproductive capacity and output early in the life cycle and a diapause-like profile later in the life cycle. Likewise, the expression levels of *mfe*, a gene encoding the final enzyme in the biosynthesis of JH and associated with fertility (Jedlicka et al., 2016) was down-regulated in late compared to early eclosing workers. Additionally, *hex1, a* gene related to diapause preparation (Denlinger, 2002; Hunt et al., 2007; Zhou et al., 2006), was up-regulated in workers eclosing later in the life cycle.

However, not all genes followed the expected trend. *Vg*, for example, is a yolk-precursor protein whose expression is expected to increase in reproductives (thus, in early-eclosing workers). However, no significant differences were found in *vg* as the colonies age. The lack of differences in *vg* could be due to the association of *vg* with aggressive behavior in bumble bees rather than solely with reproduction (Amsalem et al., 2014). The lack of differences in the expression levels of the other three genes (*foxo*, *hsp70*, and *menin*) may be explained in the fact that RNA expression do not always track the protein levels, and some genes show higher individual variability in their expression than others (in a way that our sample size may have not captured). Diapause related genes are also sometimes characterized with a complex expression pattern. *Foxo*, for example, while down-regulated in *B. terrestris* queens during diapause, is actually up-regulated in queens prior to entering diapause and right after diapause (Amsalem et al., 2015a). Additionally, *menin*, a regulator of heat shock proteins and a general stress response (Zhao and Jones, 2012), was significantly down-regulated in *B. terrestris* queens prior to entering diapause compared with queens already in diapause (Amsalem et al., 2015a).

With this in mind, we suggest that worker physiology throughout colony development matches the cascade of seasonal events occurring in the wild. Workers are expected to invest in their own reproduction when it is likely to be successful and invest in a diapause-like physiology (likely an attempt to survive the winter) when reproduction is unlikely to be successful. This explanation is fully supported by the physiological data but only partially supported by the gene expression results.

### 4. Physiological shift in workers are innate to the queen: queens lay eggs of different quality as they age

A final explanation we propose is that the observed shift in worker physiology results from the age of the queen. As social insect queens age, their reproductive output may shift to maximize their own fitness or due to stressors associated with aging. Bumble bee queens control their reproduction to delay production of gynes until later in the colony cycle, thereby reducing the time available for worker reproduction (Alaux et al., 2005). Studies examining the factors regulating gyne production have shown through cross-fostering experiments that the production of gynes can be accelerated by placing older queens (post-competition) in young colonies, suggesting that this event is controlled by the queen (Alaux et al., 2005; Alaux et al., 2006). *B. terrestris* queens also decrease in fecundity following the switch point in the colony cycle (i.e., the transition of queen to laying male eggs), reducing the frequency of egg-laying events as well as the number of eggs laid per cell (Almond et al., 2019). Queen age-induced shifts in offspring number and quality has also been identified in honey bees *(A. mellifera caucasica)* where colonies containing two-year old queens had 50% lower levels of brood production, reduced frames of adult bees and reduced brood quality compared to colonies headed by queens younger than one-year old (Akyol et al., 2007). Furthermore, mortality of larvae increased with queen age, as do the size of the eggs produced (Al-Lawati and Bienefeld, 2009).

Overall, while these studies suggest that offspring physiology may shift as the queen ages, the function of producing diapause-like workers later in the season in bumble bees is not immediately clear since the shifts we observed in worker physiology over the colony’s development do not necessarily indicate a reduction in the quality of the worker. In fact, with the exception of week 13, workers in older colonies had an increased size and mass and contained higher levels of fat body lipids, which could be interpreted as signs of increased worker quality. With that in mind, it should be noted that honey bee (*A. mellifera*) queens in smaller colonies or experiencing periods of nutritional stress were shown to produce larger eggs, likely as a way to increase the chance of their survival (Amiri et al., 2020). Although we did not control for the age of queen, our control of the environmental conditions and worker demography likely reduced the probability that the queen was simply responding to the colony environment but cannot preclude the possibility that the physiological shift in workers is due to increased queen age.

## Conclusions

Bumble bee workers were demonstrated to express a physiological signature corresponding to the age of the colony with a shift towards a diapause-like profile in late-eclosing workers. These findings may indicate that worker physiology is innate, but whether the signature is innate to workers, the colony or the queen warrants further study.

## Acknowledgments

We thank members of the Amsalem laboratory and Professor Abraham Hefetz for helpful discussions and critical reading of earlier drafts of the manuscript. We further thank two reviewers whose comments largely improved the manuscript.

