## Supplemental Information for "Shift in worker physiology and gene expression pattern from reproductive to diapause-like with colony age in the bumble bee *Bombus impatiens*"

**Table S1*:*** Information on development and size of colonies used in this study. The number of workers sampled from different colonies for each of the different analyses is indicated in parentheses.

| **Colony** | **Sample size** | **Gynes Produced** | **Males Produced** | **# Workers Produced** |
|---|---|---|---|---|
| **1** | Cold tolerance (435), ovaries, size and mass (231) | Never | Week 13 | 674 |
| **2** | Cold tolerance (485), ovaries, size and mass (241) | Never | Week 13 | 759 |
| **3** | Cold tolerance (233), lipids (26) | Week 6 (n=59) | Week 8 | 233 |
| **4** | Cold tolerance (752), lipids (33) | Never | Week 7 | 752 |
| **5 (wild)** | Cold tolerance (136), lipids (14) | Week 8 (n=30) | Week 12 | 230 |
| **6 (wild)** | Cold tolerance (375), lipids (17) | Never | Week 13 | 444 |
| **7** | Egg laying (118) | Week 10 (n=57) | Week 9 | 150 |
| **8** | Egg laying (124) | Week 9 (n=75) | Week 9 | 650 |
| **9** | Egg laying (144) | Week 10 (n=175) | Week 10 | 500 |
| **10** | Egg laying (58), Gene expression (2) | Week 8 (n=94) | Unknown | Unknown |
| **11** | Gene expression (3) | Unknown | Unknown | Unknown |
| **12** | Gene expression (3) | Unknown | Unknown | Unknown |
| **13** | Gene expression (3) | Unknown | Unknown | Unknown |
| **14** | Gene expression (3) | Unknown | Unknown | Unknown |
| **15** | Gene expression (3) | Unknown | Unknown | Unknown |

**Table S2.** The correlation between the numbers of days workers survived in 3-5°C and worker's age, body mass and oocyte size. In two of the colonies, workers were tagged and their age upon mortality was known. Body mass and ovaries were measured in workers after their death and tested against the individual survival data of workers.

| **Analysis** | **Individual Age** | **Body mass** | **Ovary activation** |
|---|---|---|---|
| **1** | *Not examined* | r=0.13, n=322, p=0.02 | r=0.05, n=278, p=0.32 |
| **2** | *Not examined* | r=0.00, n=344, p=0.93 | r=0.04, n=308, p=0.44 |
| **3** | r= -0.14, n=232, p=0.03 | r=0.22, n=233, p<0.001 | r=0.03, n=28, p=0.85 |
| **4** | r=-0.21, n=743, p<0.001 | r=0.00, n=752, p=0.58 | r=-0.02, n=43, p=0.86 |
| **5** | *Not examined* | r=0.06, n=136, p=0.48 | r=0.167, n=51, p=0.24 |
| **6** | *Not examined* | r=0.04, n=376, p=0.43 | r=0.07 n=58, p=0.62 |

**Table S3.** List of genes examined in this study, their accession numbers, and primer sequences

| Gene | Accession Numbers | Forward primer | Reverse Primer |
|---|---|---|---|
| Arginine kinase (HKG) | XM_012391881.2 | GTTGGTAGGGCAGAAGGTCA | AGGTCTACCGTCGTCTGGTG |
| Phospholipase A2 (HKG) | XM_003491149.3 | CATTTCCGCAAGTGGTAGGT | GGTCACACCGAAACCAGATT |
| Forkhead transcription factor (*foxo*) | XM_024366034.1<br>XR_002946280.1<br>XR_001102115.2<br>XR_001102114.2<br>XM_012382794.2 | CTCCCATCAATTGTCCCATC | CAACAACAATCGCAAACAGG |
| Hexamerin1 (*hex1*) | XM_012390757.2 | GAATTGCCAAATCGAGAGGA | TCATTCAATTGCCACCCCCA |
| Heat shock protein 70 (*hsp70*) | XM_003491915.3<br>XM_012389299.2 | TCATTGTCTGGTCGCAGGTC | ACCTATGGATGAAGAAAGGCGT |
| Menin (*menin*) | XM_003488543.3 | AACCGATCACTGGCAACCAT | GATCACGCCTGGGTGGTTTA |
| Methyl farneosoate epoxidase (*mfe*) | XM_003484680.3 | CAGCCGCCAATATGATACCT | GATCACGCCTGGGTGGTTTA |
| Phosphoenolpyruvate carboxykinase (*pepck)* | XM_012381009.2 | GTTGGTAGGGCAGAAGGTCA | GCTCTACGCGACCATCTCTC |
| Vitellogenin (*vg*) | XM_003492229 | CAGCCGCCAATATGATACCT | CCCTCCGTTCGAAGTGATAA |
